# Multiplexed transcriptional repression identifies a network of bactericidal interactions between mycobacterial respiratory complexes

**DOI:** 10.1101/2021.09.18.460886

**Authors:** MB McNeil, HW. Ryburn, J. Tirados, CY. Cheung, GM. Cook

## Abstract

*Mycobacterium tuberculosis* remains a leading cause of infectious disease morbidity and mortality for which new drug combination therapies are needed. Combinations of respiratory inhibitors can have synergistic or synthetic lethal interactions suggesting that regimens with multiple bioenergetic inhibitors will drastically shorten treatment times. However, realizing this potential is hampered by a lack of on-target inhibitors and a poor understanding of which inhibitor combinations have the strongest interactions. To overcome these limitations, we have used CRISPR interference (CRISPRi) to characterize the consequences of transcriptionally inhibiting individual respiratory complexes and identify bioenergetic complexes that when simultaneously inhibited result in cell death. In this study, we identified known and novel synthetic lethal interactions and demonstrate how the engineering of CRISPRi-guide sequences can be used to further explore networks of interacting gene pairs. These results provide fundamental insights into the functions of and interactions between bioenergetic complexes and the utility of CRISPRi in designing drug combinations.

## Introduction

*Mycobacterium tuberculosis* remains a significant health problem^1^. Although a curable disease requiring a combination of four drugs for six months, drug-resistant strains of *M. tuberculosis* require long treatment times with often highly toxic drugs. In addition to new drugs, rational approaches to the design of combination therapies are needed to shorten the length of treatment and prevent the emergence of drug resistance.

To generate sufficient energy to support aerobic growth, mycobacterial respiration couples the oxidation of electron donors from organic carbon catabolism (including NADH, succinate and malate) to the reduction of oxygen as a terminal electron acceptor^2,3^. Mycobacteria encode several primary dehydrogenases that can donate electrons to the electron transport chain (ETC) via the electron carrier menaquinone^2,3^. This includes two classes of NADH:menaquinone oxidoreductases that couple the oxidation of NADH to the reduction of menaquinone i.e. a proton-pumping type I NADH dehydrogenase (NDH-1) and a non-proton pumping type II NADH dehydrogenase (NDH-2); two succinate dehydrogenase enzymes that couple the oxidation of succinate to the reduction of menaquinone i.e. SDH-1 and SDH-2 and a malate:quinone oxidoreductase (MQO) that couples the oxidation of malate to the reduction of menaquinone^2-4^. Under aerobic conditions, a supercomplex of cytochrome *bc1* (Complex III; *qcrBCD*) and a *aa*_*3*_-type cytochrome *c* oxidase (Complex IV; *ctaBCDE*) couples the transfer of electrons from reduced menaquinol to cytochrome *c* to proton translocation and the generation of a proton motive force (PMF)^5,6^. An alternative cytochrome *bd*-type menaquinol oxidase (*cydAB*) functions as a purported higher affinity terminal oxidase under low oxygen conditions^7^. The generated proton gradient is used to drive ATP synthesis via the F_1_F_o_-ATP synthase (*atpBEFHAGDC)*.

Therapeutic targeting of mycobacterial respiration holds significant clinical promise, as highlighted by the FDA approval of the inhibitors bedaquiline (inhibitor of F_1_F_o_-ATP synthase)^8,9^, delamanid^10^ and the clinical testing of Q203 (inhibitor of Complex III)^11,12^. Combinations of respiratory inhibitors can also have synergistic or synthetic lethal interactions. For example, genetic or chemical inhibition of the terminal oxidase, cytochrome *bd* has synthetic lethal interaction in *M. tuberculosis* with the bacteriostatic inhibitor Q203 resulting in rapid cell death both *in vitro* and in murine infection models^7,13^.

New regimens with multiple respiratory inhibitors have the potential to drastically shorten treatment times. However, realising this potential requires novel on-target inhibitors for different respiratory components and an understanding of which combinations of inhibitors have the strongest interactions in a clinical setting.

To address this lack in knowledge we have utilised a combination of mycobacterial CRISPR interference (CRISPRi) and phenotypic assays of bacterial growth and viability to identify respiratory complexes that when simultaneously inhibited result in cell death^14,15^. These results demonstrate that the simultaneous transcriptional inhibition of the supercomplex III-IV (i.e. *qcrB*) in combination with *cydB* or *ndh*2 produces a synthetically lethal interaction that results in cell death. However, the strength of synthetic lethal interactions is influenced by the specific gene pair being targeted and mycobacterial metabolism. We also demonstrate the bactericidal consequences of inhibiting *ndh*2 or *atpE* and how sgRNA engineering to produce hypomorphic bacteriostatic sgRNAs can further resolve networks of interacting gene pairs. These results provide fundamental insights into the functional interactions between respiratory complexes, the clinical promise of targeting these processes and the general utility of CRISPRi in designing new combination therapies.

## Results

### Transcriptional repression of respiratory complexes in *M. smegmatis* has both bacteriostatic and bactericidal consequences

Using *Mycobacterium smegmatis* as physiological model for mycobacterial respiration and ATP synthesis, we investigated the consequences of transcriptionally inhibiting genes involved in mycobacterial respiration. Transcriptional repression was achieved using mycobacterial CRISPRi that uses 20-25 nucleotide sequences with complementarity to the non-template strand of target genes, termed sgRNA, to guide a deactivated Cas9 nuclease from *Streptococcus thermophilus* CRISPR1 (dCas9) to bind target genes^14-16^. Target-bound sgRNA-dCas9 impedes the progression of RNA polymerase, thereby inhibiting transcription. Both dCas9 and the sgRNA are expressed from a single integrative plasmid under the control of anhydrotetracycline (ATc)-inducible promoters^14^. Two sgRNAs per target gene were individually cloned into the CRISPRi plasmid (i.e. pLJR962). Consequences of transcriptional repression on bacterial growth and viability were determined when grown in HdeB-minimal media using succinate as a sole, non-fermentable, carbon source in a 96-well plate format. Targeting *nuoD* (i.e. complex I), *sdhA1* (i.e SDH1, complex II), *cydB* (i.e. terminal oxidase) or *menD* (i.e. menaquione biosynthesis) had no effect on cell growth or viability (Fig 1A-B). A single sgRNA targeting *menD* (i.e. *menD*_b) impaired growth, but had no effect on CFU/ml (Fig 1A and B). With the exception of *menD*, these results are consistent with previously published deletion mutants and high-throughput transposon mutagenesis, confirming that these genes are dispensable for growth under the tested conditions^17,18^. Transcriptional inhibition of *ndh2, sdhA2* (i.e. SDH2, Complex II), *mqo, qcrB* (i.e. Complex III), *ctaC* (i.e. complex IV), *atpE* or *atpB* (ATP synthase) resulted in impaired bacterial growth (Fig 1A-B).

**Figure 1.**
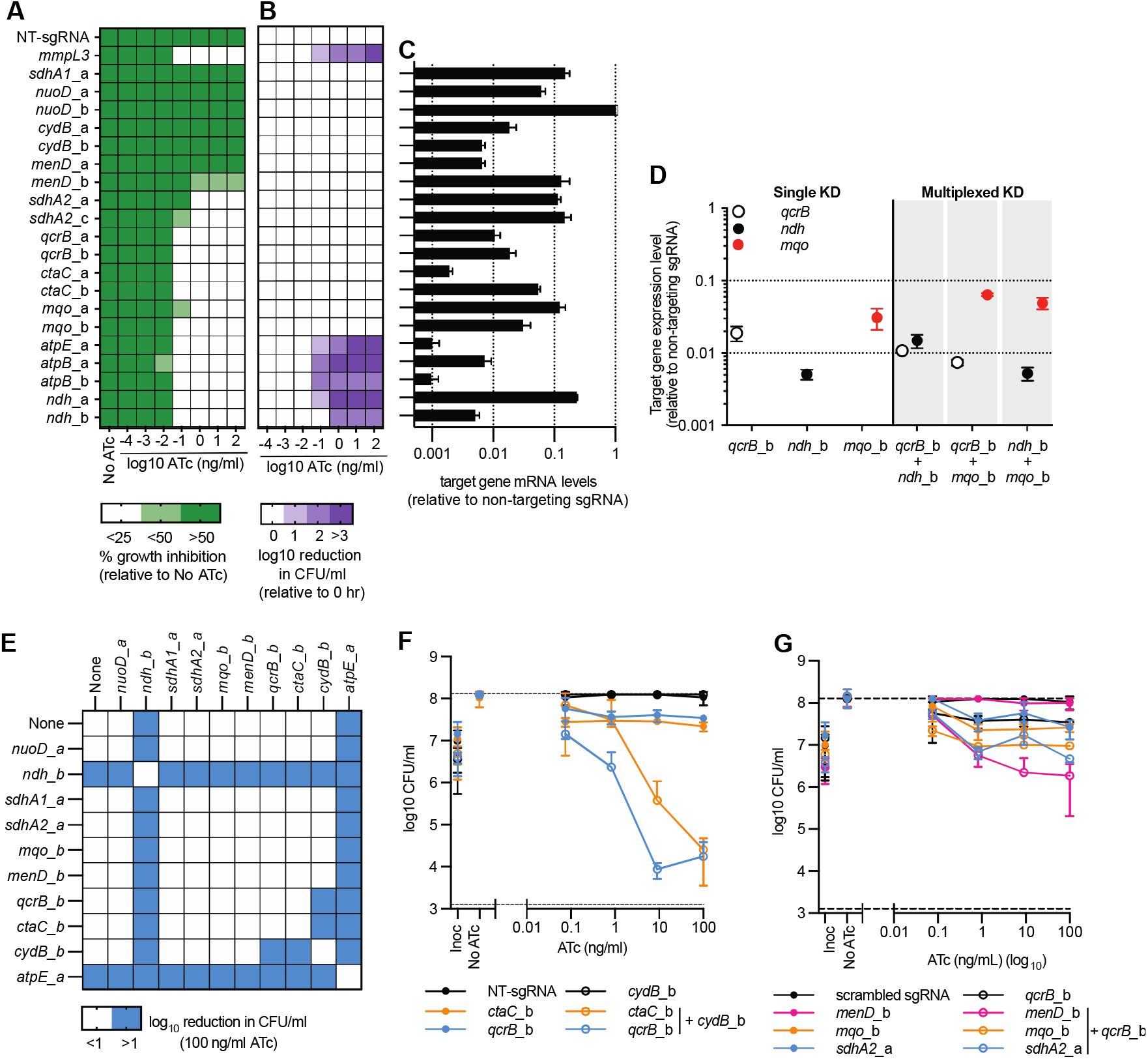
Individual and simultaneous transcriptional repression of respiratory complexes in *M. smegmatis*: (A) Growth of *M. smegmatis* expressing sgRNAs from a starting OD_600_ of 0.005 in 96 well plates in a total volume of 100 µl with differing levels of ATc. Results are mean ± standard deviation of four replicates, expressed relative to a no-ATc control. (B) Viability of *M. smegmatis* expressing sgRNAs as described for (A). Results are mean ± standard deviation of four replicates. For A and B, strains expressing either non-targeting (NT) or mmpL3 targeting sgRNA are used as a negative and positive-bactericidal control respectively. (C) mRNA levels of target genes following CRISPRi induction with 100 ng/ml when grown in LBT. mRNA levels are expressed relative to a non-targeting sgRNA. Results are mean ± standard deviation of three technical replicates. The a and b labels represent the a and b guides targeting each genetic target. (D) Transcriptional analysis of *qcrB, ndh* and *mqo* in either single or multiplexed knockdown strains. Results of targeted genes are expressed relative to a non-targeting control in the presence of 100 ng/ml ATc and are the mean ± SD of three technical replicates. (E) Log10 reduction (0 – 26 hrs) in CFU/ml for single and multiplex strains in the presence of 100 ng/ml ATc. Blue shading represents a gene pair that when simultaneously repressed have at least a one-log reduction in CFU/ml. sgRNAs paired with “none” denote results from strains that express a single sgRNA. (F-G) CFU/ml plots of selected single and multiplexed KD against increasing concentrations of ATc. A strain expressing a non-targeting (NT) sgRNA is used as a negative control. (F) Expression of sgRNAs *qcrB*_b and *ctaC* alone or in combination with sgRNAs *cydB*_b. (G) Expression of sgRNAs *menD*_b, *mqo_b* and *sdhA2*_a alone or in combination with sgRNA *qcrB*_b. For F and G, results are the mean ± SD of at least four biological replicates.

The transcriptional inhibition of *sdhA2, mqo, qcrB* and *ctaC* had bacteriostatic effects on cell viability with no reduction in detectable CFU (Figure 1B). The transcriptional inhibition of *ndh, atpB* and *atpE* had bactericidal consequences on cell viability, with >1 log reduction in CFU/ml (Figure 1B). For *atpB* and *atpE* this is consistent with our previously published work^16^. In contrast to *M. tuberculosis, M. smegmatis* has only a single gene encoding NDH-2, yet the bactericidal phenotype of inhibiting *ndh2* in this study is consistent with the CRISPRi targeting of *ndh* in *M. tuberculosis*^19^. Transcriptional analysis demonstrated that all sgRNAs, with the exception of *nuoD*_b, repressed target gene expression at least 10-fold relative to a non-targeting sgRNA control, with an absence of correlation between transcriptional repression and phenotype (Figure 1C). This suggests that variation in CRISRPi transcriptional repression does not influence the observed phenotype. In conclusion, CRISRPi is able to identify essential genes involved in mycobacterial respiration that have either a bacteriostatic or bactericidal consequence on cell viability.

### Simultaneous inhibition of mycobacterial respiratory complexes identifies interacting gene pairs

To identify synergistic or synthetically lethal interactions between mycobacterial respiratory complexes we constructed multiplexed CRISPRi plasmids that simultaneously repress the expression of two target genes. Transcriptional analysis of multiplexed strains demonstrated that target genes are repressed to comparable levels as observed in the single KD strains (Figure 1D). Phenotypic analysis demonstrated that all multiplexed combinations with at least a single essential gene repressed mycobacterial growth (Figure S1). Analysis of cell viability demonstrated that the simultaneous transcriptional repression of *qcrB* or *ctaC* with *cydB* resulted in bacterial killing (Figure 1E and F), whilst repression of individual complexes had a bacteriostatic or non-inhibitory phenotype (Figure 1E and F). Whilst there were no detectable reductions in CFU/mL, multiplexed strains of *qcrB* or *ctaC* with *sdhA2, menD* or *mqo* exhibited a significant growth defect that required at least five days of incubation before visible colonies could be counted, in contrast to the typical three days of incubation required of other strains. These multiplexed interactions also had an additive or synergistic interaction with a stronger bacteriostatic phenotype than the single sgRNAs alone (i.e. the single knockdowns had a 0.5-1 log increase in CFU/ml) (Figure 1G). The bactericidal phenotypes of *ndh* and *atpE* prevented the observation of interactions with other respiratory complexes (Figure 1E and S2A and B). In conclusion, multiplex CRISPRi identifies interactions between mycobacterial respiratory components.

### Mutagenesis identifies suboptimal sgRNAs targeting *ndh* and *atpE*

The bactericidal phenotypes of *ndh* and *atpE* prevented the detection of genetic interactions (Figure 1E and S2A and B). We hypothesized that suboptimal sgRNAs with a growth inhibitory but non-lethal phenotype would allow for the investigation of interactions between *ndh*/*atpE* and other respiratory complexes. To identify suboptimal sgRNAs we (I) mutated every second nucleotide in the sgRNAs *ndh*_b and *atpE*_a and (II) selected sgRNAs with lower predicted PAM scores targeting *ndh* and *atpB/E*. Phenotypic analysis demonstrated that mis-matched and alternative PAM targeting sgRNAs were active and retained their inhibitory effects on bacterial growth (Figure S3A-D). The majority of the mis-matched *ndh*_b sgRNAs abolished the *ndh*_b bactericidal phenotype and instead had a bacteriostatic phenotype with no reduction in CFU/ml (Figure 2A-B). Mis-match at position 11 of *ndh*_b retained the bactericidal activity (Figure 2A-B). Conversely, the majority of mis-matched *atpE*_a sgRNAs retained the bactericidal phenotype of *atpE*_a (Figure 2D-E). The one exception being a mis-match at position three, that had a bacteriostatic phenotype with no reduction in CFU/ml (Figure 2D-E). Similarly, the majority of sgRNAs with weaker PAM scores resulted in a loss of bactericidal phenotype when targeting *ndh*, but retained their bactericidal phenotype when targeting *atpE* or *atpB* (Figure S3E-F).

**Figure 2.**
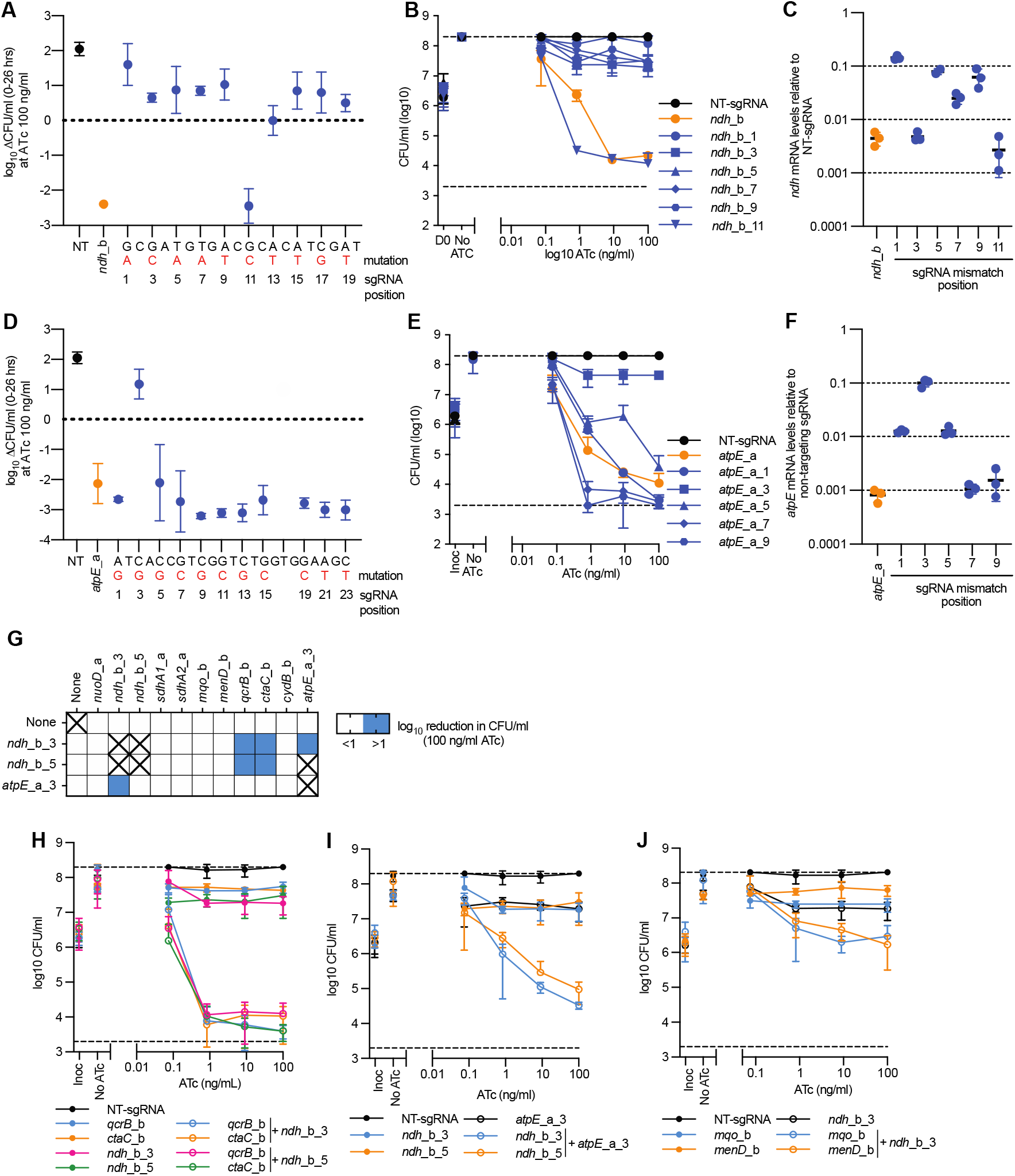
Mutagenesis of sgRNAs targeting *ndh* and *atpE* identifies suboptimal sgRNAs with unique synthetic lethal interactions: (A) log_10_ Reduction in CFU/ml (CFU/ml at 0 hrs -26hrs) in the presence of 100 ng/ml ATc for strains expressing either a non-targeting (NT), *ndh_b* or a *ndh_b* mismatched sgRNA. Mismatch base pair is highlighted in red, with the sgRNA position noted below. (B) CFU/ml plots for strains expressing either a non-targeting (NT), *ndh_b* or selected *ndh_b* mismatched sgRNA against increasing concentrations of ATc. Mismatched sgRNAs are noted by the position of their mutation in the parental sgRNA. (C) mRNA levels of *ndh_b* or selected *ndh_b* mismatched sgRNAs following CRISPRi induction with 100 ng/ml. mRNA levels are expressed relative to a non-targeting sgRNA. Results are mean ± standard deviation of three technical replicates. (D) log_10_ Reduction in CFU/ml (CFU/ml at 0 hrs -26hrs) in the presence of 100 ng/ml ATc for strains expressing either a non-targeting (NT), *atpE_a* or a *atpE_a* mismatched sgRNA. Mismatch base pair is highlighted in red, with the sgRNA position noted below. (E) CFU/ml plots for strains expressing either a non-targeting (NT), *atpE_a* or selected *atpE_a* mismatched sgRNA against increasing concentrations of ATc. Mismatched sgRNAs are noted by the position of their mutation in the parental sgRNA. (F) mRNA levels of *atpE_a* or selected *atpE_a* mismatched sgRNAs following CRISPRi induction with 100 ng/ml. mRNA levels are expressed relative to a non-targeting sgRNA. Results are mean ± standard deviation of three technical replicates. All phenotypic results (A, B, D and E) results are the mean ± SD of at least four biological replicates. (G) Log10 reduction (0 – 26 hrs) in CFU/ml for single and multiplex strains in the presence of 100 ng/ml ATc. Blue shading represents a gene pair that when simultaneously repressed have at least a one-log reduction in CFU/ml. sgRNAs paired with “none” denote results from strains that express a single sgRNA. Combinations marked with “X” were not tested. (H-J) CFU/ml plots of selected single and multiplexed KD against increasing concentrations of ATc. A strain expressing a non-targeting (NT) sgRNA is used as a negative control. (H) Expression of sgRNAs *qcrB*_b and *ctaC* alone or in combination with sgRNAs *ndh*_b_3 or *ndh*_b_5. (I) Expression of sgRNA *atpE*_a_3 alone or in combination with sgRNAs *ndh*_b_3 or *ndh*_b_5. (J) Expression of sgRNAs *mqo*_b and *menD*_b alone or in combination with sgRNA *ndh*_b_3. For H-J, results are the mean ± SD of at least four biological replicates.

Transcriptional profiling of suboptimal sgRNAs demonstrated that for the *ndh* mis-matches the majority of bacteriostatic sgRNAs had reduced levels of repression compared to the bactericidal parental sgRNA or *ndh*_b_11 (Figure 2C). Interestingly, the bacteriostatic *ndh*_b_3 had a comparable level of repression to the parental sgRNA and *ndh*_b_11 (Figure 2C). For the *atpE* mis-matches, the single bacteriostatic sgRNA repressed *atpE* expression by only 10-fold, whilst bactericidal mismatched sgRNAs repressed *atpE* expression by at least 100-fold and the parental sgRNA repressed expression by approximately 1000-fold (Figure 2F). In conclusion, mutations in sgRNA targeting sequences or weaker predicted PAM sequences reduces the level of transcriptional repression and can generated a hypomorphic phenotype.

### Suboptimal sgRNAs targeting *ndh* and *atpE* identify interacting gene pairs

To identify interactions with *atpE* and *ndh*, the suboptimal bacteriostatic sgRNAs *atpE*_a_3, *ndh*_b_3 and *ndh*_b_5 were multiplexed into CRISPRi plasmids with other respiratory components. Phenotypic analysis of these interactions demonstrated that suboptimal *ndh* targeting sgRNAs had synthetic lethal interactions with *qcrB* and *ctaC* (Figure 2G and H). Interestingly, the hypomorphic *ndh* and *atpE* targeting sgRNA also had a, albeit weaker, synthetically lethal interaction between themselves, achieving an approximately 1 log reduction in CFU/ml at the maximum ATc concentration (Figure 2G and I). The simultaneous repression of *menD* or *mqo* with the transcriptionally stronger *ndh* hypomorphic sgRNA (i.e. mis-match at position three, but not five) had an additive or synergistic interaction with a stronger bacteriostatic phenotype than the single sgRNAs alone (Figure 2J). In conclusion, the construction of suboptimal sgRNA with bacteriostatic phenotypes allows for the identification of interactions associated with the inhibition of *ndh* and *atpE*.

### Rate and magnitude of killing by simultaneous target inhibition is influenced by gene pair and metabolic state

To validate synthetic lethal interactions and to investigate possible variations in the rate or magnitude of killing, time kill experiments were performed. Time kill experiments reproduced the synthetic lethal interactions between *qcrB* and *cydB* and *qcrB* with the *ndh* hypomorphs, in addition to the weaker interaction between the *ndh* and *atpE* hypmorphs (Figure 3A-B). The synthetic lethal interaction between *qcrB* and *ndh* hypomorphs had a greater than two-log reduction within 14 hrs, whilst the *qcrB+cydB* interaction was slower and reached a one- and two-log reduction in CFU/ml at 20 and 26 hrs respectively (Figure 3A-B). The interaction between the *ndh* and *atpE* hypomorph was the weakest and reached a one log reduction in CFU/ml at 26 hrs (Figure 3A-B)

**Figure 3.**
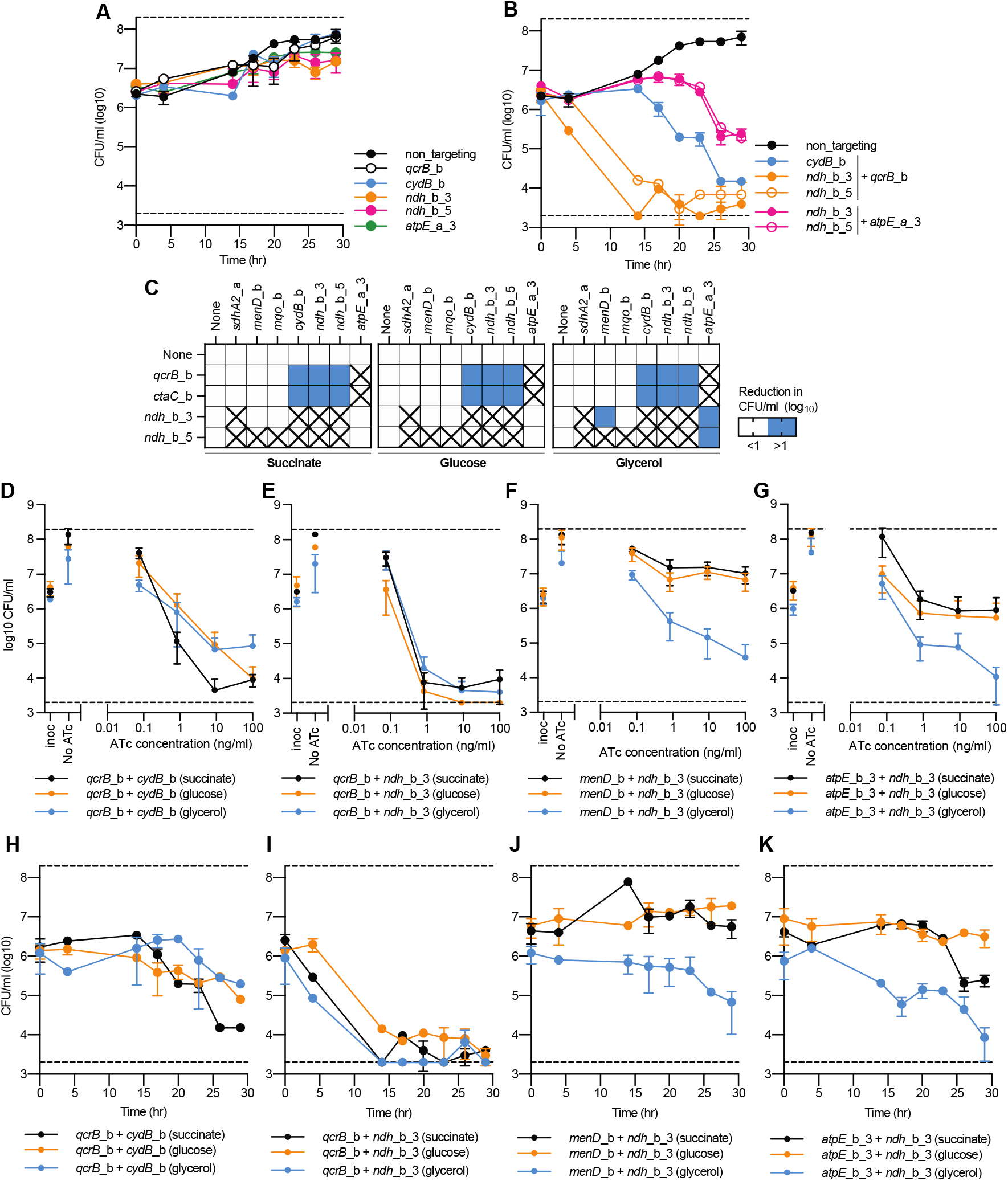
Rate and magnitude of killing by synthetic lethal interactions is influence by gene pair and growth on alternative carbon sources: (A-B) Time kill assays of either (A) single or (B) multiplexed gene pairs when grown in the presence of 100 ng/ml ATc with succinate as a sole carbon source. CFUs were taken at stated time points. For A-B, data is the mean ± SD of biological duplicates from a representative experiment. (C) Log10 reduction (0 – 26 hrs) in CFU/ml for single and multiplex strains in the presence of 100 ng/ml ATc. Blue shading represents a gene pair that when simultaneously repressed have at least a 1 log greater reduction in CFU/ml. sgRNAs paired with “none” denote results from strains that express a single sgRNA. Boxes marked with an “X”, denoted gene pairs that were not analysed on alternative carbon sources because they failed to show a synergistic or synthetic lethal interaction when previously tested (see Figure 1 and 2). (D-G) CFU/ml plots of selected single and multiplexed KD against increasing concentrations of ATc. A strain expressing a non-targeting (NT) sgRNA is used as a negative control. Black, orange and blue lines represent results when grown in either succinate, glucose, or glycerol respectively. (D-G) Reduction in CFU/ml for *M. smegmatis* strains expressing the sgRNA (D-E) *qcrB*_b in combination with sgRNAs (D) *cydB*_b or (E) *ndh*_b_3, or (F-G) expression of the sgRNA *ndh*_b_3 in combination with (F) *menD_*b or (G) *atpE_*a_3. For D-G, results are the mean ± SD of at least four biological replicates. (H-K) Kill curves for *M. smegmatis* strains expressing the sgRNA (H-I) *qcrB*_b in combination with sgRNAs (D) *cydB*_b or (E) *ndh*_b_3, or (F-G) expression of the sgRNA *ndh*_b_3 in combination with (F) *menD_*b or (G) *atpE_*a_3. For H-K, results are the mean ± SD two biological replicates from a representative experiment. Black, orange and blue lines represent results when grown in either succinate, glucose, or glycerol respectively. For H-K, the succinate kill curve is the same as presented in panel 3B.

Our experimental set up utilized succinate as a sole carbon non-fermentable source, which forces mycobacterial respiration to use the TCA cycle and ETC (i.e. oxidative phosphorylation) to generate ATP. We sought to determine whether synthetic lethal interactions would be rescued or amplified when grown on alternative carbon sources such as glucose or glycerol (i.e. fermentable carbon sources) that allow ATP to be generated via glycolysis and provide opportunities for metabolic rerouting to overcome a dysfunctional TCA cycle^20-22^. The synthetic lethal interaction between *qcrB*-*cydB* and *qcrB*-*ndh*_b_3 was conserved on all carbon sources (Figure 3C-E). The magnitude of killing by the *qcrB*-*cydB* interaction was greater when grown in succinate as a sole carbon source, with a 1 log increase in killing at 10 ng/ml Atc (Figure 3D). There was no variation in the magnitude of killing by the *qcrB*-*ndh*_b_3 interaction across different carbon sources (Figure 3E). Interestingly, the *ndh*_b_3 interaction with either *menD* or *atpE*_b_3 became bactericidal when grown with glycerol as a sole carbon, with a >1 log reduction in CFU/ml at 100 ng/ml ATc (Figure 3C and F-G). Time kill experiments validated the consistency of killing by the *qcrB*-*ndh*_b_3 interaction across different carbon sources, with a greater than two-log reduction in CFU/ml reach by 14 hrs (Fig 3I). The *qcrB-cydB* interaction whilst present across all carbon sources was stronger when grown in succinate with a two-log reduction in CFU/ml at 26 hrs, compared to a one-log reduction when grown on glucose or glycerol (Fig 3H). Consistent with ATc titration results, the *ndh*_b_3 interaction with *menD* was bactericidal when grown on glycerol as a sole carbon source with a one-log reduction at 30 hrs, compared to no reduction in CFU/ml when grown on succinate or glucose (Fig 3J). Similarly, the *ndh*_b_3 interaction with *atpE*_b_3 had the greatest level of killing when grown on glycerol as a sole carbon source with a >two-log reduction in CFU/ml at 30 hrs compared to the one-log or less than one-log reduction when grown on succinate or glucose respectively (Fig 3K). In conclusion, the rate and magnitude of killing resulting from the simultaneous inhibition of respiratory components is influenced by both the genes pairs that are targeted and mycobacterial metabolic state.

## Discussion

New drugs and drug regimens are urgently needed to combat *M. tuberculosis*. Mycobacterial respiration and metabolism have emerged as a promising area for the development of novel drugs and regimens. This is highlighted by the approval of the ATP synthase inhibitor BDQ and the cytochrome-*bc*_1_ inhibitor, Q203, being advanced into clinical trials^9,12^. There is also evidence that combinations of respiratory inhibitors have the potential for synergistic or synthetically lethal interactions with sterilizing activity, supporting the development of novel regimens involving multiple respiratory inhibitors^7,23,24^. Herein we have used mycobacterial CRISPRi to identify the cellular consequences of individually and simultaneously inhibiting the major respiratory components of oxidative phosphorylation in mycobacteria. We demonstrate that in *M. smegmatis*, the majority of respiratory components are essential and there is a network of interactions centered around the inhibition of the CIII-CIV supercomplex.

Inhibition of *menD* produced a growth defect, but was not identified as an essential gene under the tested conditions. This is in contrast to previous TnSeq experiments in *M. tuberculosis* and CRISPRiseq studies in *M. smegmatis* where the *menD* is designated as an essential gene. These discrepancies maybe a combination of (i) differences in experimental set up (growth in liquid vs selection on solid media), (ii) CRISPRi being unable to achieve sufficient levels of inhibition to produce a phenotype or (iii) because following repression of *menD* residual menaquinone remains available and is sufficient to buffer the genetic depletion and support bacterial growth for a period of time^19,25^. Consistent with previous results, *sdh1* and *nuoD* are non-essential ^4,17,26,27^. The inhibition of *sdh2, qcrB, ctaC* and *mqo* produced an inhibitory phenotype with bacteriostatic consequences on cell viability.

For *qcrB*, this is consistent with previous chemical inhibitors including Q203 and TB47^11,24,28^. The inhibition of *ndh* and *atpE* and *atpB* had an inhibitory phenotype with bactericidal consequences. These results for *atpE/B* and *ndh2* are consistent with our recent studies in *M. tuberculosis*^*16,19*^. It is important to note, there are genetic redundancies in *M. tuberculosis* with it encoding two copies of *ndh-2* (i.e. *ndh* and *ndhA*) and in contrast to CRISPRi mediated transcriptional inhibition, previous targeted deletion strategies have demonstrated that in *M. tuberculosis* only when both copies of *ndh* (i.e. *ndh* and *ndhA*) are deleted in the presence of fatty acids is an essential phenotype observed^26,27^. Resolving the discrepancies between these genetic approaches requires additional work, but suggests that chemical inhibitors of NDH-2 should have a bactericidal phenotype as has been observed for clofazimine and quinolinequinones^29,30^.

Simultaneous transcriptional inhibition of *cydB* and *qcrB*, which had non-essential and bacteriostatic phenotypes respectively, resulted in cell death. This synthetic lethal interaction is consistent with previous chemical-genetic and chemical-chemical studies in mycobacteria^7,13^. This also validates the use CRISPRi to identify favourable interactions between possible drug targets. Multiplexed interactions also demonstrated that *qcrB* and *ctaC*, together which form the CIII-CIV supercomplex in mycobacteria^5,6^, have additive or potentially synergistic interactions with *sdhA2, mqo* and *menD*. Multiplexed interaction analysis of hypomorphic sgRNAs also identified a synthetic lethal interaction between *qcrB*/*ctaC* and *ndh* as well as a synthetic lethal interaction between the inhibition of *atpE* and *ndh*. The rate and magnitude of killing by synthetic lethal interactions varied greatly, with *ndh+qcrB* being the strongest, followed by *qcrB+cydB*, and then the *ndh+atpE* hypomorph. Growth on alternative carbon sources also influenced synthetic lethal interactions, with the interaction between *ndh* + *menD* or *atpE* being greater when grown on glycerol, whilst the interaction between *qcrB+cydB* was stronger when grown with succinate as a sole carbon source. The retention of the *qcrB+cydB* interaction across diverse carbon sources is consistent with previous chemical-genetic studies in *M. tuberculosis*^*31*^. Combined, this work has significantly expanded the network of synthetic lethal interactions between mycobacterial respiratory components, demonstrated that the rate and magnitude of synthetic lethal interactions is highly variable and highlighted the potential for efficacious combination therapies that consist of multiple respiratory inhibitors.

This study also has wider implications on the understanding of CRISPR applications and the generation of suboptimal hypomorphic sgRNA sequences to probe interactions between genetic targets. Previous studies of *Streptococcus pyogenes* Cas9 demonstrated that as little as five bp similarity in the seed region (i.e. proximal to PAM sequence) is sufficient to facilitate sgRNA binding with disruption of this region perturbing sgRNA targeting^32^. Conversely, in this study the transcriptional effects of sgRNA mutations in *S. thermophilus* Cas9 was variable, with mutations in the *atpE* sgRNA showing a positional effect (i.e. mutations closer to PAM had a weaker transcriptional repression), whilst mutations in the *ndh2* sgRNA lacked a positional effect. Suboptimal sgRNAs targeting *atpE* suggest that between 10-100 fold transcriptional repression is sufficient to prevent growth but not kill, whilst above 100 fold repression results in bacterial killing. Conversely, for suboptimal sgRNAs targeting *ndh* even with no perturbations in transcriptional repression cellular outcomes can be changed. This suggests that other factors, such as dCas9 retention on target sequences, are likely to be important factors for when considering the efficacy of mutated sgRNAs. With a limited dataset of only two sgRNAs the results of this current study cannot provide any general design principles for constructing hypomorphic sgRNAs when using this mycobacterial CRISPRi platform. However, the contrasting phenotypic consequences for suboptimal guides targeting *ndh* (i.e. majority bacteriostatic) and *atpE* (i.e. majority bactericidal) suggest that even with predictable effects of mutagenesis on sgRNA efficacy, as are available for *S. pyogenes* dCas9^33^, the effects on cellular phenotype are completely dependent on the gene being targeted and the relative level of transcriptional inhibition needed to produce a phenotype.

Several of our genetic interactions have been previously reported using chemical inhibitors. This includes (1) the synthetic lethal interaction between *qcrB* (i.e. Q203) and *cydB* deletion^7^, and the synergistic killing between (2) *ndh* (i.e. clofazimine) and *qcrB* (i.e. Q203 and IMP)^24^, (3) *menA* and *qcrB* (i.e. IMP)^23^ and (4) *ndh* (i.e. clofazimine) and *atpE* (i.e. bedaquiline)^24^. The respiratory inhibitor C10 with a currently undefined mechanism of action is also synthetically lethal with Q203, suggesting that *qcrB* may have synthetic lethal interactions with other respiratory components outside of those investigated in this current study^34^. For the interaction between clofazimine with either Q203 or bedaquiline, it has been proposed that Q203 and bedaquiline both deplete ATP levels whilst increasing carbon metabolism and TCA cycle activity^24^. The proposed increase in TCA cycle activity results in an accumulation of reducing equivalents that cause reductive stress and potentiate clofazimines ability to transfer electrons from NDH-2 to oxygen resulting in rapid production of reactive oxygen species and bacterial killing^24^. Whether this potential mechanism of simultaneously starving mycobacteria of energy to limit their ability to utilise antioxidant defences in the face of increasing oxidative stress is universal to all of our observed synthetic lethal interactions requires further investigation. It is also worth noting that previous studies have identified an interaction between chemical inhibition of menaquinone biosynthesis and chemical inhibition of either *atpE* (i.e. bedaquiline) or *ndh* (i.e. clofazimine)^23,35^. The lack of a genetic interaction between *menD* + *atpE* and the *menD* + *ndh* interaction only being observed when grown on glycerol, maybe be a combination of *menD* transcriptional inhibition being insufficient or because of metabolic buffering that following repression of *menD*, residual menaquinone remains and is sufficient to support bacterial growth for a period of time^19^. Alternatively, this lack of chemical validation in some cases may be due to potential off target effects of chemical inhibitors.

Combined, the results from this current study have significantly expanded the network of synthetic lethal interactions between mycobacterial respiratory components, the majority of which involve the inhibition of the cytochrome-*bc*_1_-*aa*_3_ supercomplex. Previous work highlighting the synthetic lethality of the interaction between the terminal oxidase *cydB* and *qcrB*, has placed significant emphasis on the identification of cytochrome bd oxidase inhibitors in order to realise the full potential of Q203. These results expand the potential suite of drug targets that could be combined with Q203. In conclusion, this work supports the development of novel respiratory inhibitor combinations for the treatment of mycobacterial infections, should energize the discovery of small molecule inhibitors targeting other respiratory components and highlights the power of CRISPRi in identifying unique combination therapies.

## Material and Methods

### Bacterial strains and growth conditions

*Escherichia coli* MC1061 and *M. smegmatis* mc^2^155 were grown in luria broth (LB) media or LB agar (1.5%) at 37°C and shaking at 200 rpm when required. For growth of *M. smegmatis* in liquid LB media tween-80 was added at 0.05% (LBT). *M. smegmatis* phenotypic experiments were performed in a minimal Hartmans de bont (HdeB) media (2g (NH_4_)SO_4_, 1.08 g KH_2_PO_4_ and 2.99 g Na_2_HPO_4_ with 10 ml 100x trace metals per L^36^) supplemented with tyloxapol (0.05%) and either succinate (30 mM), glucose (20 mM) or glycerol (0.2%) as a sole carbon source. When necessary media was supplemented with Kanamycin at 50 µg/ml for *E. coil* and 25 µg/ml for *M. smegmatis*.

### Construction and transformation of CRISPRi plasmids

A 20-25 bp sequence downstream of permissible PAM sequences targeting the non-template strand of target genes of interest were identified^14^. Target sequences were ordered as oligos with GGGA and AAAC overhangs respectively (Table S1), and cloned into pJLR962 using BsmB1 and golden gate cloning as previously described^15^. Plasmids were cloned into *E. coli* and validated with sanger sequencing. CRISPRi plasmids expressing multiple sgRNA were constructed by cloning the promoter-sgRNA-dCas9 scaffold from single expression plasmids using the primer pair MMO120 + MMO121 (Table S2). PCR products were purified and cloned into target single sgRNA expression plasmids using Sap1 and golden gate cloning, following the protocol described in Table S3. Plasmids were cloned into *E. coli* and validated with sanger sequencing. Confirmed single and double sgRNA CRISPRi expression plasmids are listed in Table S1 and 4 respectively. Plasmids were electroporated into *M. smegmatis* mc^2^155 following previously established protcols^15,16^.

### Quantification of transcriptional knockdowns

To quantify the level of transcriptional inhibition, *M. smegmatis* strains expressing sgRNA were grown from a starting OD_600_ of 0.1 in HdeB-Succinate with 100 ng/ml ATc. Cultures were grown for 16 hrs at 37°C with shaking at 200 rpm. RNA was extracted, cDNA was synthesized and qPCR reactions were performed as previously described protocol^15^. The removal of DNA was confirmed using 1 µl of extracted RNA and the primer combination MMO202 + MMO203. qPCR reactions were performed using the primer pairs listed in Table S2. Signals were normalized to the housekeeping *sigA* transcript and quantified by the 2^ΔΔCt^ method^37^. Error bars are the standard deviation of three technical replicates.

### CRISPRi phenotypic assessment of essentiality and viability

To determine the consequences of targeted gene repression on bacterial growth, phenotypic assays were performed as follows (Figure S4). *M. smegmatis* strains containing sgRNA expression plasmids were initially grown overnight in 5 ml HdeB-succinate + Kan. Cultures were diluted to an OD_600_ of 0.01 in HdeB-succinate + Kan and 50 µl was used to inoculate a 96 well assay plate at a starting OD_600_ of 0.005. 96 well assay plates were prepared as follows. Fifty µl of HdeB-succinate + Kan was added to all wells of column 2-11 of a 96 well plate except row H. 55 µl of HdeB-succinate + Kan containing the starting concentration of ATc (i.e. 200 ng/ml ATc) was added to row H. ATc was diluted along the vertical axis of the 96 well plate, transferring 5 µl between Rows, up to Row B. Row A was used as a no ATc control. Columns 1 and 12 contained 100 µl of media as contamination and background controls. Inoculated plates were grown at 37°C with shaking at 200 rpm for 26 hrs. OD_600_ was measured using a Varioskan-LUX microplate reader. The minimal inhibitory concentration (MIC) of ATc was determined relative to the growth of the no-ATc control, using a non-linear fitting of data to the Gompertz equation^38^. Targeted genes classified as essential for growth generated an MIC for ATc, whilst non-essential genes failed to generate an MIC. To determine the consequences of targeted gene repression on bacterial viability, colony forming units were determined at 0 and 26 hrs. Briefly, at 0 hrs a 4-point ten-fold dilution of the 0.01 diluted culture was performed in PBS and 5 µl of each dilution was spotted onto to LBA. At 26 hrs, approximately 50 µl of culture was removed from the 96 well assay plate used in essentiality screens after OD_600_ was determined. A 4-point ten-fold dilution was performed in PBS and 5 µl of each dilution was spotted onto to LBA. Plates were incubated at 37°C for 3-4 days, at which points colonies were counted. Essential genes and gene pairs with bacteriostatic consequences resulted in no change in CFU/ml relative to 0 hrs, whilst bactericidal consequences produced at least a 1 log reduction in CFU/ml relative to 26 hrs. For cultures assessing essentiality and viability with alternative carbon sources, experiments were performed as described above, with the exception that succinate was substituted for alternative carbon sources throughout and used at stated concentrations.

### Time kills assays

Time kill assays were performed by growing *M. smegmatis* strains containing sgRNA expressing plasmids overnight in 5 ml HdeB-succinate + Kan. Cultures were diluted to an OD_600_ of 0.1 in HdeB-succinate + Kan and 500 µl was used to inoculate into a final volume of 10 ml HdeB-succinate + Kan in a 100 ml flask, to give a starting OD_600_ of 0.005. Flasks were incubated at 37°C with shaking at 200 rpm. Samples were taken for OD_600_ and CFU at stated time points. OD_600_ was measured in a Jenway 6300 spectrophotometer. CFUs were determined as described above for viability assays in a 96 well plate. For time kill experiments with alternative carbon sources, experiments were performed as described above, with the exception that succinate was substituted for alternative carbon sources throughout and used at stated concentrations.

## Acknowledgments

This research was financially supported by the Maurice Wilkins Centre for Molecular Biodiscovery, the Marsden Fund (Royal Society of New Zealand) (grant number UOO1807) and the Health Research Council of New Zealand (grant number 20/459).

We have no conflicts of interest to declare.

The data that support the findings of this study are available from the corresponding author upon request.

