## Supplemental Figures for "Multiplexed transcriptional repression identifies a network of bactericidal interactions between mycobacterial respiratory complexes"

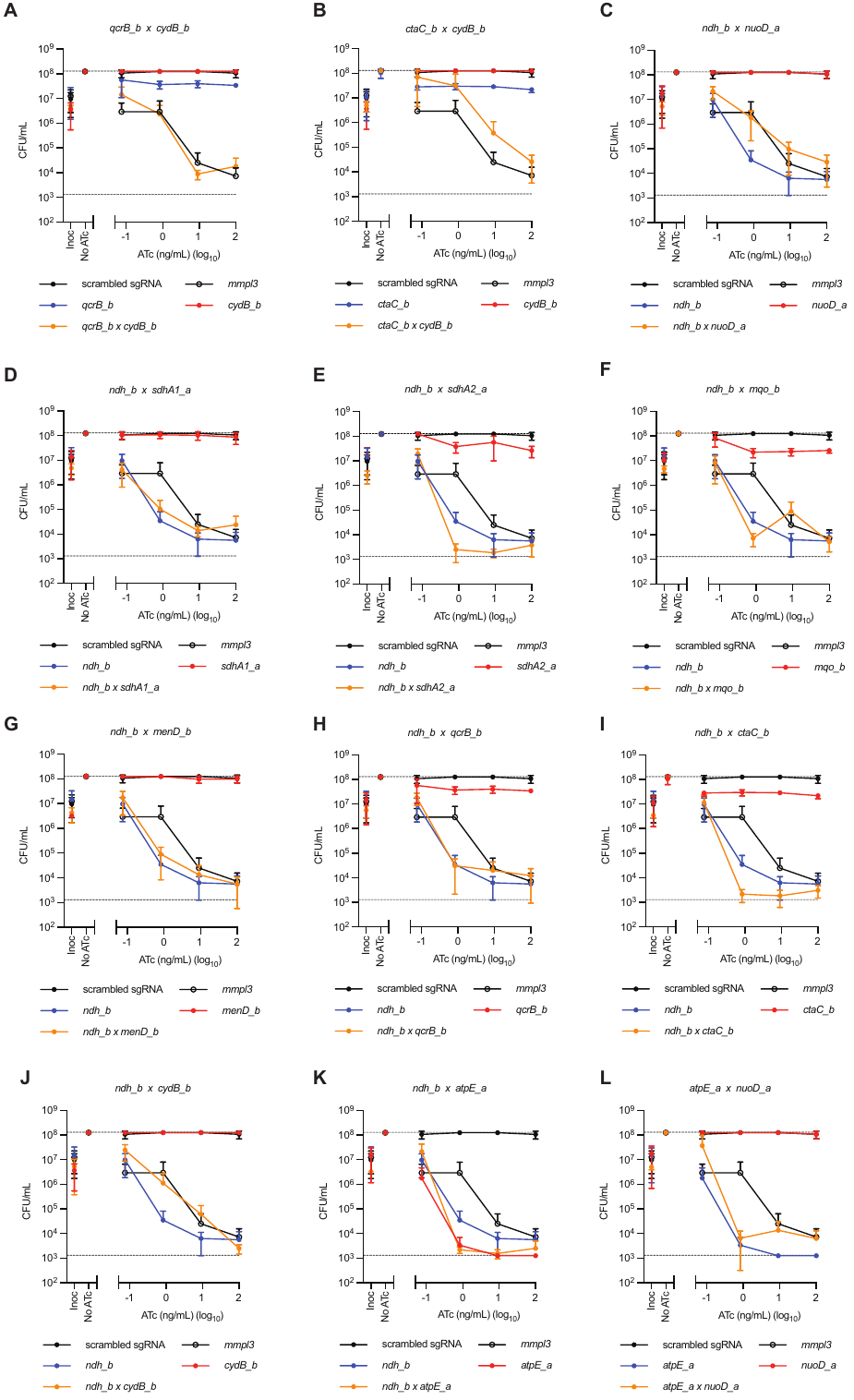

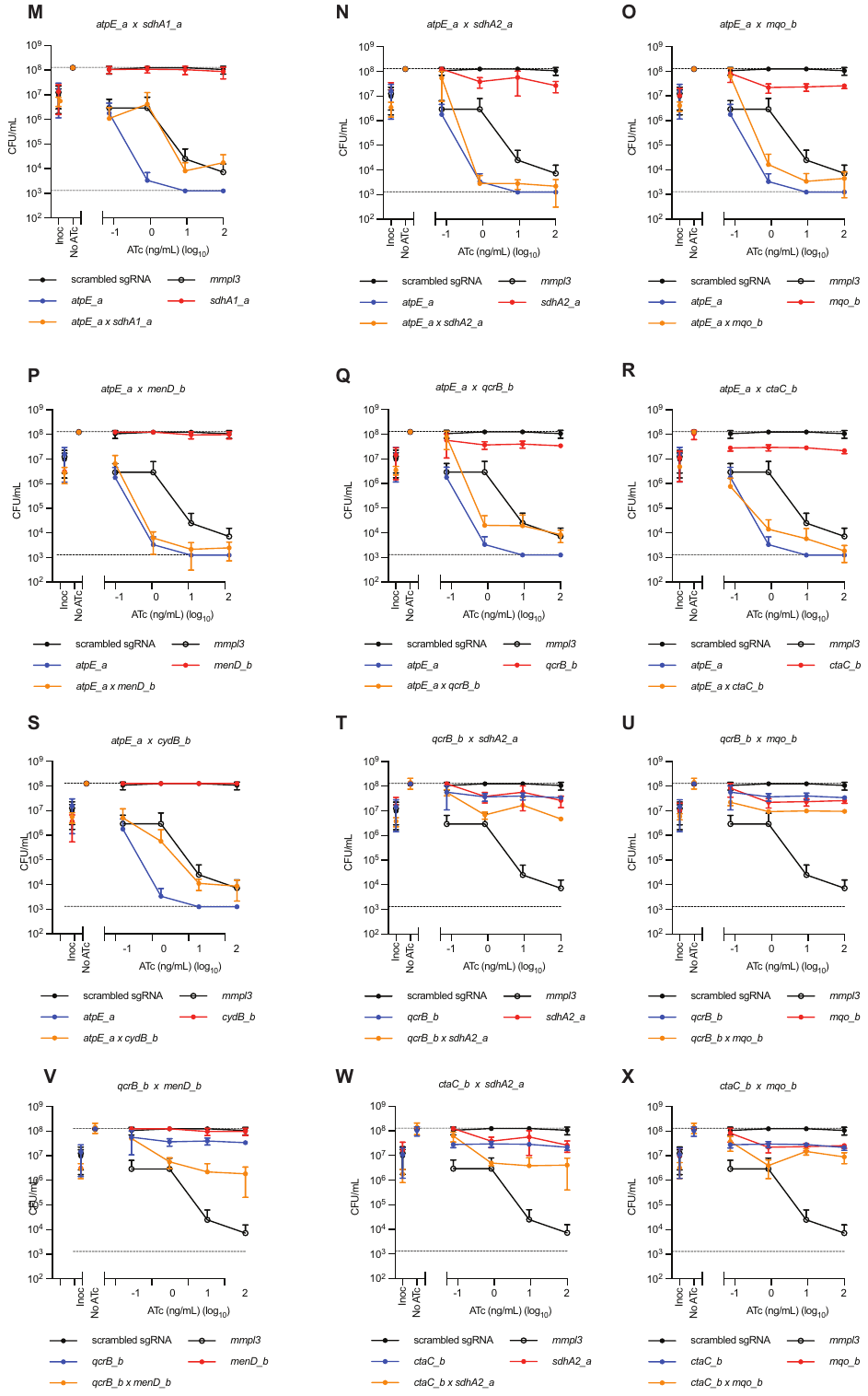

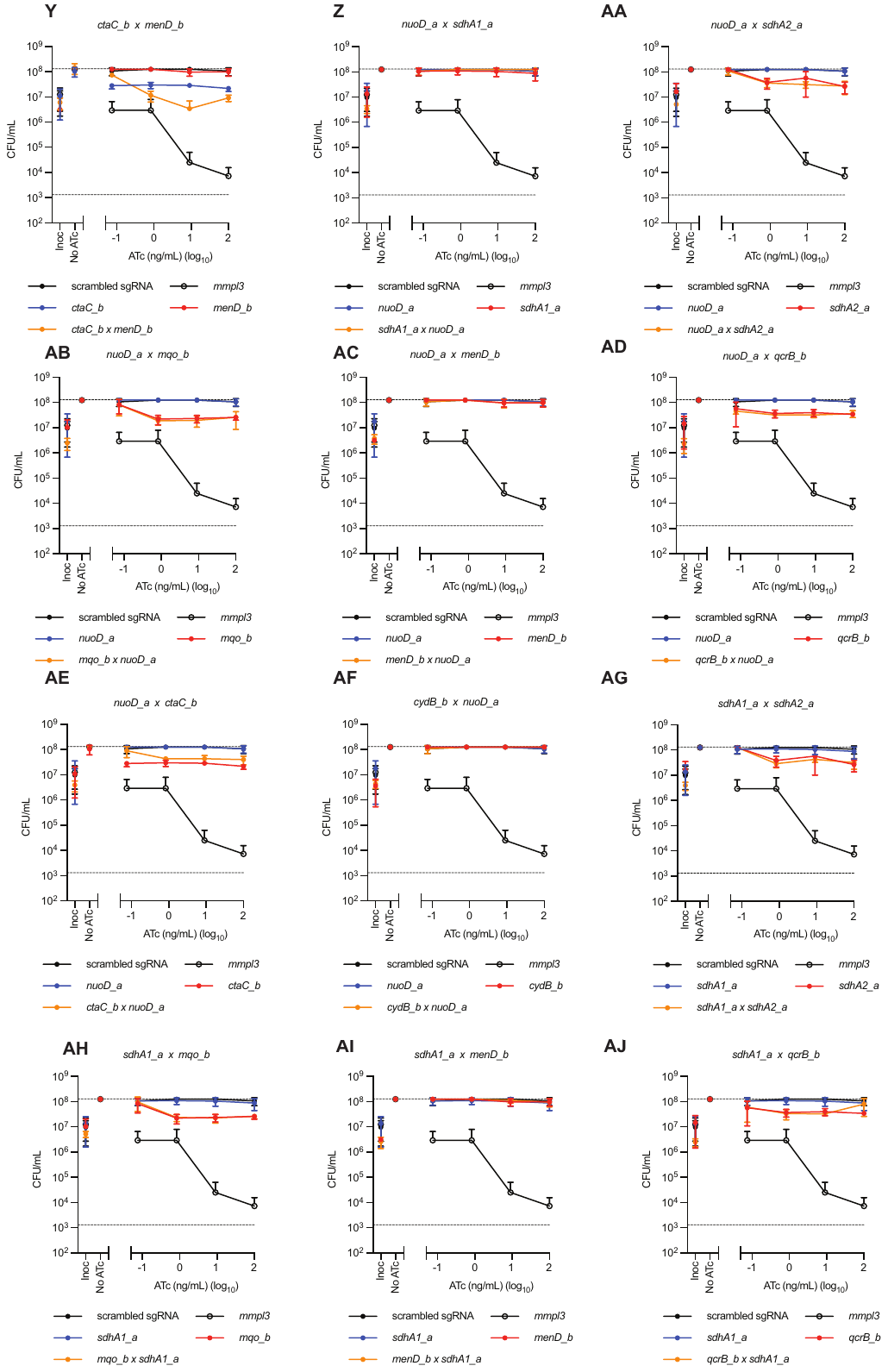

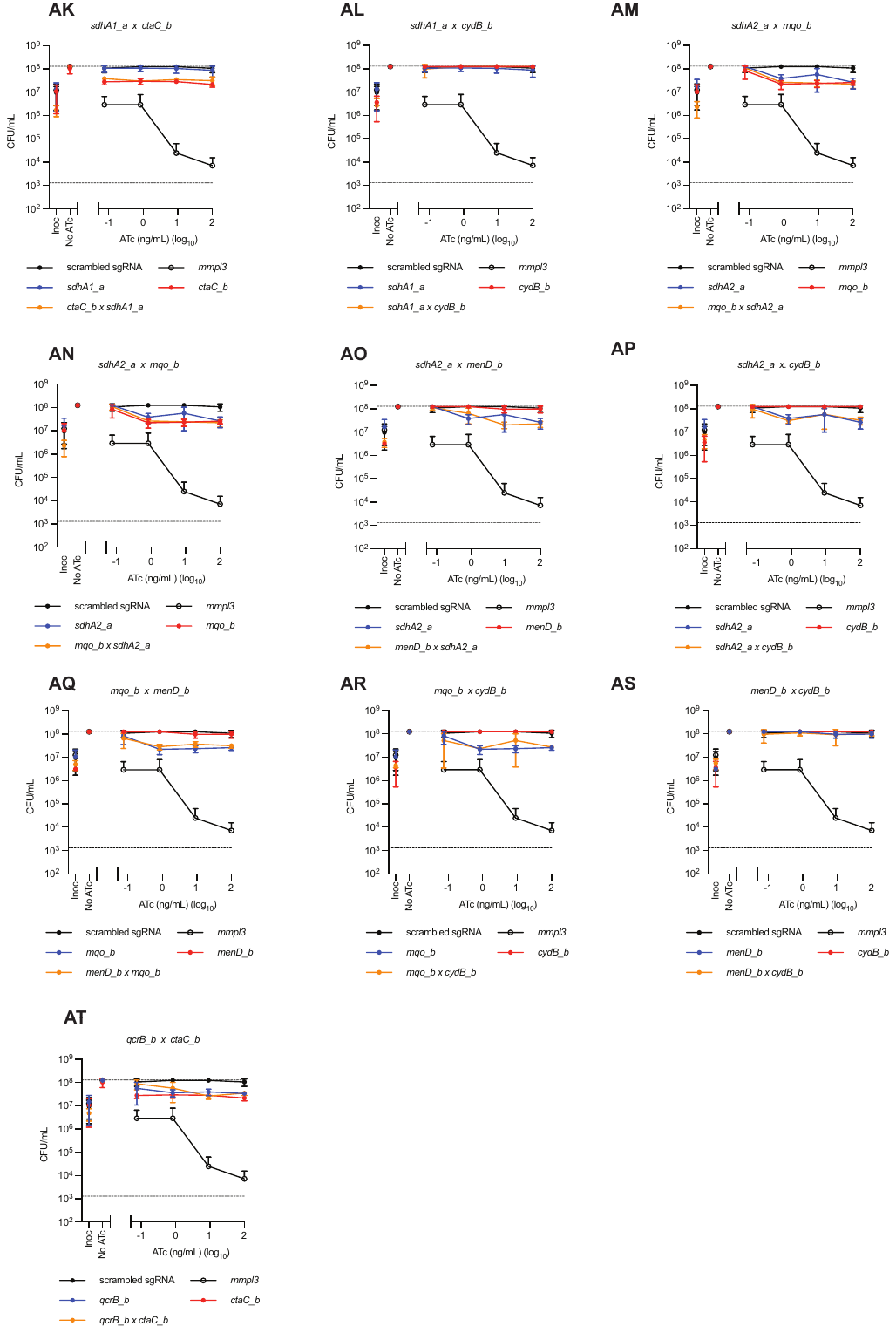

**Figure S1| Killing phenotypes of multiplexed sgRNAs targeting *ndh* and *atpE* in multiplexed combinations:** (A-AT) CFU/ml plots of *M. smegmatis* strains expressing stated single and multiplexed sgRNAs. A strain expressing a non-targeting (NT) sgRNA or sgRNA targeting *mmpL3* is used as a negative and positive control respectively*.* Results for A-AT are the mean ± SD of four biological replicates

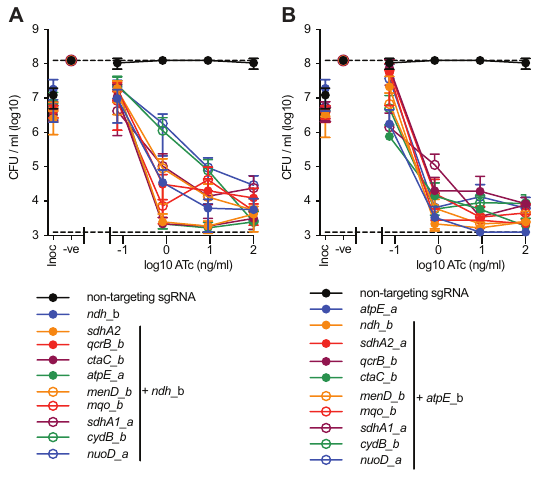

**Figure S2| Killing phenotypes of sgRNAs targeting *ndh* and *atpE* in multiplexed combinations:** (A-B) CFU/ml plots of *M. smegmatis* strains expressing (A) *ndh_b* or (B) *atpE_a* in combination with sgRNAs targeting alternative respiratory components across increasing concentrations of ATc. A strain expressing a non-targeting (NT) sgRNA is used as a negative control*.* Results for A-D are the mean ± SD of four biological replicates.

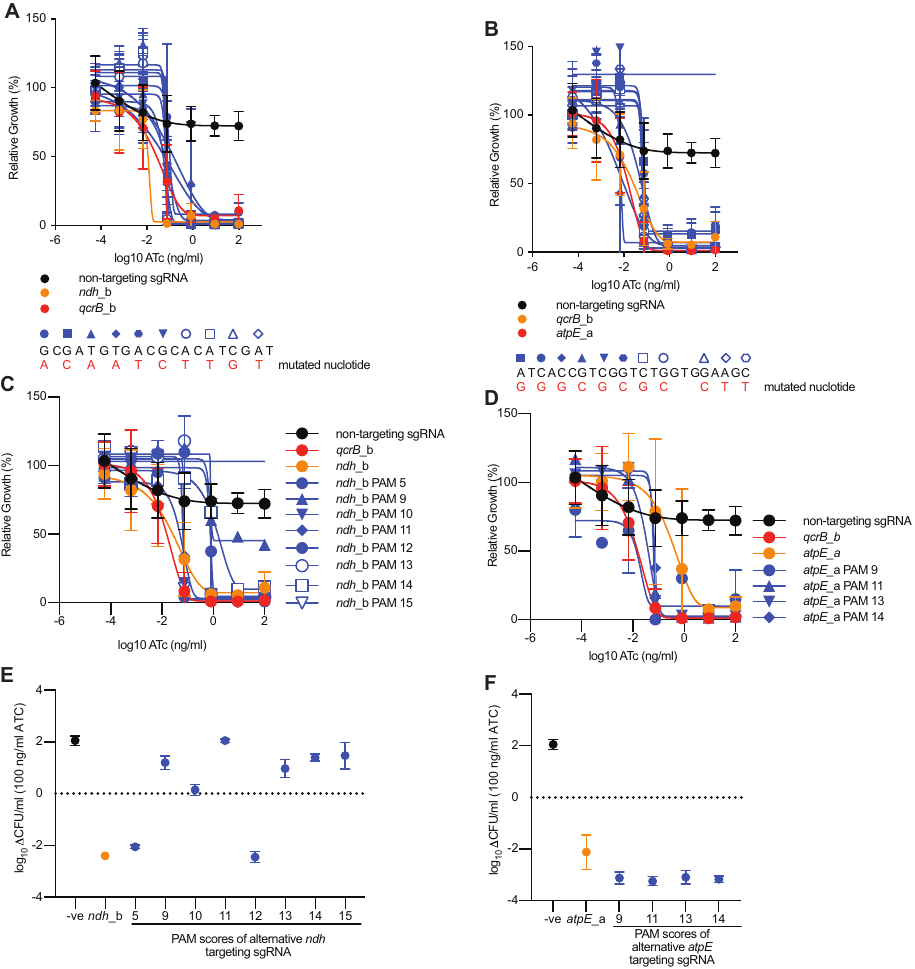

**Figure S3| Phenotypes of alternative PAM variants of sgRNAs targeting *ndh* and *atpE*:** (A-D) Growth of *M. smegmatis* strains expressing either (A) *ndh_*b mis-matched sgRNAs, (B) *atpE_*a mis-matched sgRNAs, (C) sgRNAs targeting *ndh* with weaker PAM variants or (D) sgRNAs targeting *atpE* with weaker PAM variants in the presence of increasing concentrations of ATc. Growth is expressed relative to a no ATc control. PAM score is based on previous studies that identified 15 permissible non-canonical PAM variants that retained inhibitory phenotypes^1^. (E-F) Reduction in CFU/ml (CFU/ml at 0 hrs -26hrs) in the presence of 100 ng/ml ATc. For *M. smegmatis* strains expressing sgRNAs targeting *ndh* or *atpE* with weaker PAM variants. Results for A-F are the mean ± SD of four biological replicates. For all experiments included a negative non-targeting and a parental bactericidal sgRNA (i.e. *ndh*_b or *atpE*_a)

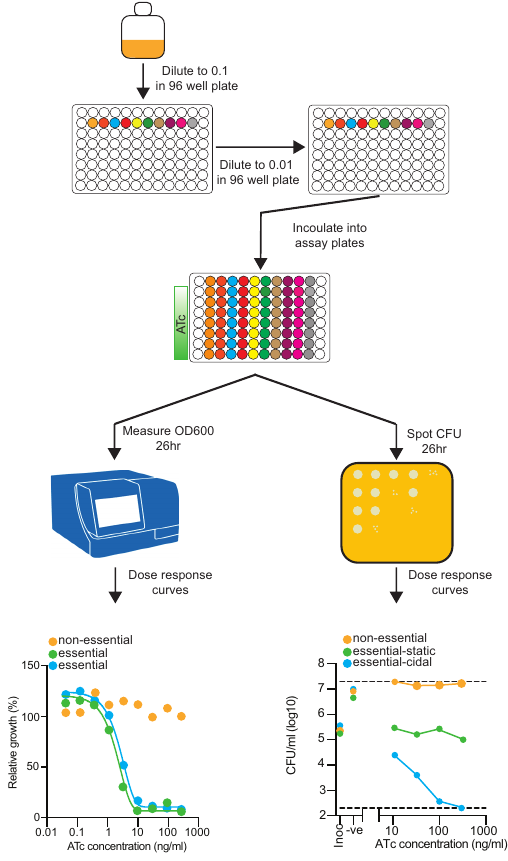

**Figure S4| Workflow for bacterial phenotyping screens of single and multiplex sgRNAs.** Work flow for phenotypic assays, with specific details in materials and methods.
