## Supplemental Tables for "Multiplexed transcriptional repression identifies a network of bactericidal interactions between mycobacterial respiratory complexes"

| **Table S1: CRISPRi plasmids used and constructed in this study** | | | | | | | | | |
| --- | --- | --- | --- | --- | --- | --- | --- | --- | --- |
| **plasmid name** | **Target-Species** | **Target-Gene** | **sgRNA name** | **Target sequence (Coding (5’-3’))** | **PAM (non coding 5’-3’, NN…...)** | **PAM Score (Rock et al)** | **sgRNA target length** | **Fwd Oligo (5-3’) (GGGA_)** | **Oligo Rev Sequence (AAAC_)** |
| pJLR962 | M. smegmatis |  | non-targeting sgRNA | CGAGACGCATTAATCGTCTCC |  |  |  | GGGAGGAGACGATTAATGCGTCTCG | AAACCGAGACGCATTAATCGTCTCC |
| pCi2 | M. smegmatis | MSMEG_0250 | *mmpL3* | GACGAGGGCAGCCAGTCTGTCGC | GTAGAAA |  |  | GGGAGCGACAGACTGGCTGCCCTCGTC | AAACGACGAGGGCAGCCAGTCTGTCGC |
| pCi3 | M. smegmatis | MSMEG_0418 | *sdhA1*_a | GCGACGAGGTGGCCCGCGCC | GGGGAAG |  |  | GGGAGGCGCGGGCCACCTCGTCGC | AAACGCGACGAGGTGGCCCGCGCC |
| pCi4 | M. smegmatis | MSMEG_1670 | *sdhA2*_a | AGGCCAAGCTGCCCGACATC | CGAGAAC | 5 | 20 | GGGAGATGTCGGGCAGCTTGGCCT | AAACAGGCCAAGCTGCCCGACATC |
| pCi21 | M. smegmatis | MSMEG_1670 | *sdhA2_*c | AGACGCTGTACCAAAACTGC | GCAGGAT | 9 | 20 | GGGAGCAGTTTTGGTACAGCGTCT | AAACAGACGCTGTACCAAAACTGC |
| pCi46 | M. smegmatis | MSMEG_2060 | *nuoD*_a | AGATCGAGGGCGAGATCATC | CCAGGAT | 9 | 20 | GGGAGATGATCTCGCCCTCGATCT | AAACAGATCGAGGGCGAGATCATC |
| pCi47 | M. smegmatis | MSMEG_2060 | *nuoD*_b | GAACGCGAGGAGATCCTGCGGGT | ACGGAAG | 4 | 23 | GGGAACCCGCAGGATCTCCTCGCGTTC | AAACGAACGCGAGGAGATCCTGCGGGT |
| pCi48 | M. smegmatis | MSMEG_3621 | *ndh*_a | CCGCTGCTCTACCAGGTGGC | CTGGAAG | 4 | 20 | GGGAGCCACCTGGTAGAGCAGCGG | AAACCCGCTGCTCTACCAGGTGGC |
| pCi49 | M. smegmatis | MSMEG_3621 | *ndh*_b | GCGATGTGACGCACATCGAT | CGAGAAG | 1 | 20 | GGGAATCGATGTGCGTCACATCGC | AAACGCGATGTGACGCACATCGAT |
| pCi50 | M. smegmatis | MSMEG_4263 | *qcrB*_a | CTGGGTGAGATCGCGCTGTAC | CAGGAAG | 4 | 21 | GGGAGTACAGCGCGATCTCACCCAG | AAACCTGGGTGAGATCGCGCTGTAC |
| pCi51 | M. smegmatis | MSMEG_4263 | *qcrB*_b | GATCCGTCGATGGCACACGT | GAAGAAC | 5 | 20 | GGGAACGTGTGCCATCGACGGATC | AAACGATCCGTCGATGGCACACGT |
| pCi52 | M. smegmatis | MSMEG_4268 | *ctaC*_a | GATTGCGTCGTTCGCCGTGGGT | ACGGAAC | 11 | 22 | GGGAACCCACGGCGAACGACGCAATC | AAACGATTGCGTCGTTCGCCGTGGGT |
| pCi53 | M. smegmatis | MSMEG_4268 | *ctaC*_b | TTCACCGTCGTCGTGCAGGAAC | GTAGAAC | 5 | 22 | GGGAGTTCCTGCACGACGACGGTGAA | AAACTTCACCGTCGTCGTGCAGGAAC |
| pCi54 | M. smegmatis | MSMEG_4941 | *atpE*_a | ATCACCGTCGGTCTGGTGGAAGC | GAAGAAC | 5 | 23 | GGGAGCTTCCACCAGACCGACGGTGAT | AAACATCACCGTCGGTCTGGTGGAAGC |
| pCi55 | M. smegmatis | MSMEG_4942 | *atpB*_a | CCACTGCCGTCACCGCCGTGAT | CGAGAAT | 2 | 22 | GGGAATCACGGCGGTGACGGCAGTGG | AAACCCACTGCCGTCACCGCCGTGAT |
| pCi56 | M. smegmatis | MSMEG_4942 | *atpB*_b | CTGCGGGCCAAGGTCACCTC | GTAGAAG | 1 | 20 | GGGAGAGGTGACCTTGGCCCGCAG | AAACCTGCGGGCCAAGGTCACCTC |
| pCi59 | M. smegmatis | MSMEG_2613 | *mqo*_a | GCCCACGCCGTCGAGAACGGT | CCAGAAC | 5 | 21 | GGGAACCGTTCTCGACGGCGTGGGC | AAACGCCCACGCCGTCGAGAACGGT |
| pCi60 | M. smegmatis | MSMEG_2613 | *mq*o_b | AACCCTGTGCCGCATGTGAGT | GAGGAAG | 4 | 21 | GGGAACTCACATGCGGCACAGGGTT | AAACAACCCTGTGCCGCATGTGAGT |
| pCi65 | M. smegmatis | MSMEG_3232 | *cydB*_a | CTGCTGGAGGGCTTCGACTT | GAAGAAG | 1 | 20 | GGGAAAGTCGAAGCCCTCCAGCAG | AAACCTGCTGGAGGGCTTCGACTT |
| pCi66 | M. smegmatis | MSMEG_3232 | *cydB*_b | GGCCGTGCCGCCGAGAAACGC | GAAGAAC | 5 | 21 | GGGAGCGTTTCTCGGCGGCACGGCC | AAACGGCCGTGCCGCCGAGAAACGC |
| pCi67 | M. smegmatis | MSMEG_1109 | *menD*_a | TGATCGTGCTGAGCGCCAAC | GCGGAAC | 11 | 20 | GGGAGTTGGCGCTCAGCACGATCA | AAACTGATCGTGCTGAGCGCCAAC |
| pCi68 | M. smegmatis | MSMEG_1109 | *menD*_b | GCACGGGCGCCAACCAGACC | CGAGCAT | 7 | 20 | GGGAGGTCTGGTTGGCGCCCGTGC | AAACGCACGGGCGCCAACCAGACC |
| pCi266 | M. smegmatis | MSMEG_3621 | *ndh*_b_1MM | ACGATGTGACGCACATCGAT | CGAGAAG | 1 | 20 | GGGAATCGATGTGCGTCACATCGT | AAACACGATGTGACGCACATCGAT |
| pCi267 | M. smegmatis | MSMEG_3621 | *ndh*_b_3MM | GCCATGTGACGCACATCGAT | CGAGAAG | 1 | 20 | GGGAATCGATGTGCGTCACATGGC | AAACGCCATGTGACGCACATCGAT |
| pCi268 | M. smegmatis | MSMEG_3621 | *ndh*_b_5MM | GCGAAGTGACGCACATCGAT | CGAGAAG | 1 | 20 | GGGAATCGATGTGCGTCACTTCGC | AAACGCGAAGTGACGCACATCGAT |
| pCi269 | M. smegmatis | MSMEG_3621 | *ndh*_b_7MM | GCGATGAGACGCACATCGAT | CGAGAAG | 1 | 20 | GGGAATCGATGTGCGTCTCATCGC | AAACGCGATGAGACGCACATCGAT |
| pCi270 | M. smegmatis | MSMEG_3621 | *ndh*_b_9MM | GCGATGTGTCGCACATCGAT | CGAGAAG | 1 | 20 | GGGAATCGATGTGCGACACATCGC | AAACGCGATGTGTCGCACATCGAT |
| pCi271 | M. smegmatis | MSMEG_3621 | *ndh*_b_11MM | GCGATGTGACCCACATCGAT | CGAGAAG | 1 | 20 | GGGAATCGATGTGGGTCACATCGC | AAACGCGATGTGACCCACATCGAT |
| pCi272 | M. smegmatis | MSMEG_3621 | *ndh*_b_13MM | GCGATGTGACGCTCATCGAT | CGAGAAG | 1 | 20 | GGGAATCGATGAGCGTCACATCGC | AAACGCGATGTGACGCTCATCGAT |
| pCi273 | M. smegmatis | MSMEG_3621 | *ndh*_b_15MM | GCGATGTGACGCACTTCGAT | CGAGAAG | 1 | 20 | GGGAATCGAAGTGCGTCACATCGC | AAACGCGATGTGACGCACTTCGAT |
| pCi274 | M. smegmatis | MSMEG_3621 | *ndh*_b_17MM | GCGATGTGACGCACATGGAT | CGAGAAG | 1 | 20 | GGGAATCCATGTGCGTCACATCGC | AAACGCGATGTGACGCACATGGAT |
| pCi275 | M. smegmatis | MSMEG_3621 | *ndh*_b_19MM | GCGATGTGACGCACATCGTT | CGAGAAG | 1 | 20 | GGGAAACGATGTGCGTCACATCGC | AAACGCGATGTGACGCACATCGTT |
| pCi276 | M. smegmatis | MSMEG_3621 | *ndh*_m | CGGGACCATCCGTCGCATCGAGT | TCGGAAT | 12 | 23 | GGGAACTCGATGCGACGGATGGTCCCG | AAACCGGGACCATCCGTCGCATCGAGT |
| pCi277 | M. smegmatis | MSMEG_3621 | *ndh*_n | CGTTCACCGTCGTCGGCGCGGGC | TCAGCAA | 10 | 23 | GGGAGCCCGCGCCGACGACGGTGAACG | AAACCGTTCACCGTCGTCGGCGCGGGC |
| pCi278 | M. smegmatis | MSMEG_3621 | *ndh*_o | GCAAGCAGAAGAACGCCCAGGT | GGAGGAT | 9 | 22 | GGGAACCTGGGCGTTCTTCTGCTTGC | AAACGCAAGCAGAAGAACGCCCAGGT |
| pCi279 | M. smegmatis | MSMEG_3621 | *ndh*_p | TTCGGCAACGACCACTTCGC | GTAGGAC | 15 | 20 | GGGAGCGAAGTGGTCGTTGCCGAA | AAACTTCGGCAACGACCACTTCGC |
| pCi280 | M. smegmatis | MSMEG_3621 | *ndh*_q | GCGGTGGCCAAGGTGGGTCC | GGAGAAC | 5 | 20 | GGGAGGACCCACCTTGGCCACCGC | AAACGCGGTGGCCAAGGTGGGTCC |
| pCi281 | M. smegmatis | MSMEG_3621 | *ndh*_r | GTCACACCTACTCGACGCCC | CGAGCAG | 13 | 20 | GGGAGGGCGTCGAGTAGGTGTGAC | AAACGTCACACCTACTCGACGCCC |
| pCi282 | M. smegmatis | MSMEG_3621 | *ndh*_s | GCACCAAGCGTGGTCAGCTC | TCAGGAA | 14 | 20 | GGGAGAGCTGACCACGCTTGGTGC | AAACGCACCAAGCGTGGTCAGCTC |
| pCi283 | M. smegmatis | MSMEG_3621 | *ndh*_t | AGCACCAAGCGTGGTCAGCT | CAGGAAC | 11 | 20 | GGGAAGCTGACCACGCTTGGTGCT | AAACAGCACCAAGCGTGGTCAGCT |
| pCi284 | M. smegmatis | MSMEG_4941 | *atpE*_a_1MM | GTCACCGTCGGTCTGGTGGAAGC | GAAGAAC | 5 | 23 | GGGAGCTTCCACCAGACCGACGGTGAC | AAACGTCACCGTCGGTCTGGTGGAAGC |
| pCi285 | M. smegmatis | MSMEG_4941 | *atpE*_a_3MM | ATGACCGTCGGTCTGGTGGAAGC | GAAGAAC | 5 | 23 | GGGAGCTTCCACCAGACCGACGGTCAT | AAACATGACCGTCGGTCTGGTGGAAGC |
| pCi286 | M. smegmatis | MSMEG_4941 | *atpE*_a_5MM | ATCAGCGTCGGTCTGGTGGAAGC | GAAGAAC | 5 | 23 | GGGAGCTTCCACCAGACCGACGCTGAT | AAACATCAGCGTCGGTCTGGTGGAAGC |
| pCi287 | M. smegmatis | MSMEG_4941 | *atpE*_a_7MM | ATCACCCTCGGTCTGGTGGAAGC | GAAGAAC | 5 | 23 | GGGAGCTTCCACCAGACCGAGGGTGAT | AAACATCACCCTCGGTCTGGTGGAAGC |
| pCi288 | M. smegmatis | MSMEG_4941 | *atpE*_a_9MM | ATCACCGTGGGTCTGGTGGAAGC | GAAGAAC | 5 | 23 | GGGAGCTTCCACCAGACCCACGGTGAT | AAACATCACCGTGGGTCTGGTGGAAGC |
| pCi289 | M. smegmatis | MSMEG_4941 | *atpE*_a_11MM | ATCACCGTCGCTCTGGTGGAAGC | GAAGAAC | 5 | 23 | GGGAGCTTCCACCAGAGCGACGGTGAT | AAACATGAGCGTCGGTCTGGTGGAAGC |
| pCi290 | M. smegmatis | MSMEG_4941 | *atpE*_a_13MM | ATCACCGTCGGTGTGGTGGAAGC | GAAGAAC | 5 | 23 | GGGAGCTTCCACCACACCGACGGTGAT | AAACATCACCGTCGGTGTGGTGGAAGC |
| pCi291 | M. smegmatis | MSMEG_4941 | *atpE*_a_15MM | ATCACCGTCGGTCTCGTGGAAGC | GAAGAAC | 5 | 23 | GGGAGCTTCCACGAGACCGACGGTGAT | AAACATCACCGTCGGTCTCGTGGAAGC |
| pCi292 | M. smegmatis | MSMEG_4941 | *atpE*_a_17MM | ATCACCGTCGGTCTGGAGGAAGC | GAAGAAC | 5 | 23 | GGGAGCTTCCTCCAGACCGACGGTGAT | AAACATCACCGTCGGTCTGGAGGAAGC |
| pCi293 | M. smegmatis | MSMEG_4941 | *atpE*_a_19MM | ATCACCGTCGGTCTGGTGCAAGC | GAAGAAC | 5 | 23 | GGGAGCTTGCACCAGACCGACGGTGAT | AAACATCACCGTCGGTCTGGTGCAAGC |
| pCi294 | M. smegmatis | MSMEG_4941 | *atpE*_a_21MM | ATCACCGTCGGTCTGGTGGATGC | GAAGAAC | 5 | 23 | GGGAGCATCCACCAGACCGACGGTGAT | AAACATCACCGTCGGTCTGGTGGATGC |
| pCi295 | M. smegmatis | MSMEG_4941 | *atpE*_a_23MM | ATCACCGTCGGTCTGGTGGAAGT | GAAGAAC | 5 | 23 | GGGAACTTCCACCAGACCGACGGTGAT | AAACATCACCGTCGGTCTGGTGGAAGT |
| pCi296 | M. smegmatis | MSMEG_4942 | *atpB*_n | CGCCGATCAACATCGTCGAAGAAC | CGAGGAA | 14 | 24 | GGGAGTTCTTCGACGATGTTGATCGGCG | AAACCGCCGATCAACATCGTCGAAGAAC |
| pCi297 | M. smegmatis | MSMEG_4942 | *atpB*_o | CACGCGGCAGGCATCTGGCGT | GTAGCAG | 13 | 21 | GGGAACGCCAGATGCCTGCCGCGTG | AAACCACGCGGCAGGCATCTGGCGT |
| pCi298 | M. smegmatis | MSMEG_4942 | *atpB*_p | TGGTACATCCAGTGGTTCCC | GGGGAAC | 11 | 20 | GGGAGGGAACCACTGGATGTACCA | AAACTGGTACATCCAGTGGTTCCC |
| pCi299 | M. smegmatis | MSMEG_4942 | *atpB*_q | TCTCCAACTGGCTCGCGGTGC | TCAGGAT | 9 | 21 | GGGAGCACCGCGAGCCAGTTGGAGA | AAACTCTCCAACTGGCTCGCGGTGC |

| **Table S2: Oligos Used in This Study** | | |
| --- | --- | --- |
| **Name-MMO** | **Sequence** | **Description** |
| MMO117 | TGCGGCGCTTTTTTTTTTGAATTC | Sequencing pJR962 |
| MMO119 | CTGCGTTATCCCCTGATTCTG | Sequencing pJR962 |
| MMO120 | AATATGCTCTTCAGGATCTGACCAGGGAAAATAGCC | Fwd primer-Cloning into SapI site pJR962/5 |
| MMO121 | TTTATGCTCTTCACTGAAAAAAAAAACACCCTGCCATAAAATGAC | Rev primer-Cloning into SapI site pJR962/5 |
| MMO202 | GCCTGGCCATCATGGGTATC | Msm_gDNA_check_F1 |
| MMO203 | GGAGGATCCGGAGACCAAGC | Msm_gDNA_check_R1 |
| MMO206 | CCTCCGTCTTTTCGGCAACA | qPCR-AtpB-MSMEG4942 |
| MMO207 | GTCGAAGGTCTTCCACACGG | qPCR-AtpB-MSMEG4942 |
| MMO208 | CTGATCTCGGGTATCGCCC | qPCR-AtpE-MSMEG4941 |
| MMO209 | GCCAGGTTGATGAAGTACGC | qPCR-AtpE-MSMEG4941 |
| MMO212 | GAGTTCGTCCTGAACTCGGC | qPCR-CtaC-MSMEG4268 |
| MMO213 | TGTCCGAGTTGTTGGCCTTC | qPCR-CtaC-MSMEG4268 |
| MMO214 | GCCTGACGATCTACAACGGA | qPCR-CydB-MSMEG3232 |
| MMO215 | GAGATGCGCTTGCTGAACAC | qPCR-CydB-MSMEG3232 |
| MMO216 | TCATAGGCGACCTGACGTTC | qPCR-MenD-MSMEG1109 |
| MMO217 | TTGTCGTTGGACACCACGAT | qPCR-MenD-MSMEG1109 |
| MMO218 | GGTGGGTCTGCTCAAGTACC | qPCR-Mqo-MSMEG2613 |
| MMO219 | GCGAATTCACGAAGCGTCTC | qPCR-Mqo-MSMEG2613 |
| MMO220 | TACGCCGCGAAGATCATCAA | qPCR-Ndh-MSMEG3621 |
| MMO221 | TGTCGAAGTACTCGAACGGC | qPCR-Ndh-MSMEG3621 |
| MMO222 | CGAGCACATCGCCAAGATCA | qPCR-nuoD-MSMEG2060 |
| MMO223 | CCTTCGGTGACGAGCTTGAA | qPCR-nuoD-MSMEG2060 |
| MMO226 | TATCGGCATGGTGGTACTGC | qPCR-qcrB-MSMEG4263 |
| MMO227 | GAAGTCGCTTGATGATGCCG | qPCR-qcrB-MSMEG4263 |
| MMO228 | CGTCATGGGTGGTATCGAGG | qPCR-sdhA1-MSMEG0418 |
| MMO229 | AGATCTGACAGCGAGTTGCC | qPCR-sdhA1-MSMEG0418 |
| MMO230 | ACAACACCAACGTCATCCCC | qPCR-sdhA2-MSMEG1670 |
| MMO231 | TGATGTCCAGCAGCGAGTTG | qPCR-sdhA2-MSMEG1670 |
| MMO270 | TGTGGGACGAGGAAGAGTCC | sigA_Msmeg_Fwd qPCR primer_set 1 |
| MMO271 | CACCTCTTCTTCGGCGTTGA | sigA_Msmeg_Rev qPCR primer_set 1 |
| MMO300 | GACTCTTCCTCGTCCCACAC | sigA_Msmeg_Fwd qPCR primer_set 2 |
| MMO301 | GAAGACACCGACCTGGAACT | sigA_Msmeg_Rev qPCR primer_set 2 |

| **Table S3: Multiplex golden gate cloning protocol** | | |
| --- | --- | --- |
| **Component** | **Volume (µl)** | |
| 10× T4 Ligase Buffer | 1 | |
| Single sgRNA expression plasmid uncut (20 ng/µL) | 1.25 | |
| Cloned and purified target -promoter-sgRNA-scaffold | 2.5 | |
| Sap1 (10,000u/ml) | 0.5 | |
| T4 DNA Ligase | 0.5 | |
| mQ | 4.25 | |
| **Temperature (℃)** | **Duration** | **Cycle** |
| 37 (digestion) | 5 min | 30 |
| 16 (ligation) | 5 min |  |
| 4 | Forever | 1 |

| **Table S4: Multiplexed Plasmids Constructed in this Study.** | | | | | | |
| --- | --- | --- | --- | --- | --- | --- |
| **Plasmid name** | **Target-Species** | **pCi plasmid backbone** | **Second sgRNA** | **Amplification template for second sgRNA** | **Fwd Primer for amplification of sgRNA module** | **Rev primer for amplification of sgRNA module** |
| pCiMX40 | *M. smegmatis* | pCi49(*ndh*_b) | *sdhA2*_a | pCi4 | MMO120 | MMO121 |
| pCiMX41 | *M. smegmatis* | pCi49(*ndh*_b) | *qcrB*_b | pCi51 | MMO120 | MMO121 |
| pCiMX42 | *M. smegmatis* | pCi49(*ndh*_b) | *ctaC*_b | pCi53 | MMO120 | MMO121 |
| pCiMX43 | *M. smegmatis* | pCi49(*ndh*_b) | *atpE*_a | pCi54 | MMO120 | MMO121 |
| pCiMX46 | *M. smegmatis* | pCi4(*sdhA2*_a) | *qcrB*_b | pCi51 | MMO120 | MMO121 |
| pCiMX47 | *M. smegmatis* | pCi4(*sdhA2*_a) | *ctaC*_b | pCi53 | MMO120 | MMO121 |
| pCiMX48 | *M. smegmatis* | pCi4(*sdhA2*_a) | *atpE*_a | pCi54 | MMO120 | MMO121 |
| pCiMX49 | *M. smegmatis* | pCi51(*qcrB*_b) | *ctaC*_b | pCi53 | MMO120 | MMO121 |
| pCiMX50 | *M. smegmatis* | pCi51(*qcrB*_b) | *atpE*_a | pCi54 | MMO120 | MMO121 |
| pCiMX51 | *M. smegmatis* | pCi53(*ctaC*_b) | *atpE*_a | pCi54 | MMO120 | MMO121 |
| pCiMX52 | *M. smegmatis* | pCi68(*menD*_b) | *ndh*_b | pCi49 | MMO120 | MMO121 |
| pCiMX53 | *M. smegmatis* | pCi68(*menD*_b) | *sdhA2*_a | pCi4 | MMO120 | MMO121 |
| pCiMX54 | *M. smegmatis* | pCi68(*menD*_b) | *qcrB*_b | pCi51 | MMO120 | MMO121 |
| pCiMX55 | *M. smegmatis* | pCi68(*menD*_b) | *ctaC*_b | pCi53 | MMO120 | MMO121 |
| pCiMX56 | *M. smegmatis* | pCi68(*menD*_b) | *atpE*_a | pCi54 | MMO120 | MMO121 |
| pCiMX57 | *M. smegmatis* | pCi60(*mqo*_b) | *ndh*_b | pCi49 | MMO120 | MMO121 |
| pCiMX58 | *M. smegmatis* | pCi60(*mqo*_b) | *sdhA2*_a | pCi4 | MMO120 | MMO121 |
| pCiMX59 | *M. smegmatis* | pCi60(*mqo*_b) | *qcrB*_b | pCi51 | MMO120 | MMO121 |
| pCiMX60 | *M. smegmatis* | pCi60(*mqo*_b) | *ctaC*_b | pCi53 | MMO120 | MMO121 |
| pCiMX61 | *M. smegmatis* | pCi60(*mqo*_b) | *atpE*_a | pCi54 | MMO120 | MMO121 |
| pCiMX62 | *M. smegmatis* | pCi60(*mqo*_b) | *menD*_b | pCi68 | MMO120 | MMO121 |
| pCiMX237 | *M. smegmatis* | pCi3 (*sdhA1*_a) | *ndh*_b | pCi49 | MMO120 | MMO121 |
| pCiMX238 | *M. smegmatis* | pCi3 (*sdhA1*_a) | *sdhA2*_a | pCi4 | MMO120 | MMO121 |
| pCiMX239 | *M. smegmatis* | pCi3 (*sdhA1*_a) | *qcrB*_b | pCi51 | MMO120 | MMO121 |
| pCiMX240 | *M. smegmatis* | pCi3 (*sdhA1*_a) | *ctaC*_b | pCi53 | MMO120 | MMO121 |
| pCiMX241 | *M. smegmatis* | pCi3 (*sdhA1*_a) | *atpE*_a | pCi54 | MMO120 | MMO121 |
| pCiMX242 | *M. smegmatis* | pCi3 (*sdhA1*_a) | *menD*_b | pCi68 | MMO120 | MMO121 |
| pCiMX243 | *M. smegmatis* | pCi3 (*sdhA1*_a) | *mq*o_b | pCi60 | MMO120 | MMO121 |
| pCiMX244 | *M. smegmatis* | pCi66 (*cydB*_b) | *ndh*_b | pCi49 | MMO120 | MMO121 |
| pCiMX245 | *M. smegmatis* | pCi66 (*cydB*_b) | *sdhA2*_a | pCi4 | MMO120 | MMO121 |
| pCiMX246 | *M. smegmatis* | pCi66 (*cydB*_b) | *qcrB*_b | pCi51 | MMO120 | MMO121 |
| pCiMX247 | *M. smegmatis* | pCi66 (*cydB*_b) | *ctaC*_b | pCi53 | MMO120 | MMO121 |
| pCiMX248 | *M. smegmatis* | pCi66 (*cydB*_b) | *atpE*_a | pCi54 | MMO120 | MMO121 |
| pCiMX249 | *M. smegmatis* | pCi66 (*cydB*_b) | *menD*_b | pCi68 | MMO120 | MMO121 |
| pCiMX250 | *M. smegmatis* | pCi66 (*cydB*_b) | *mq*o_b | pCi60 | MMO120 | MMO121 |
| pCiMX251 | *M. smegmatis* | pCi66 (*cydB*_b) | *sdhA1*_a | pCi3 | MMO120 | MMO121 |
| pCiMX252 | *M. smegmatis* | pCi46 (*nuoD*_a) | *ndh*_b | pCi49 | MMO120 | MMO121 |
| pCiMX253 | *M. smegmatis* | pCi46 (*nuoD*_a) | *sdhA2*_a | pCi4 | MMO120 | MMO121 |
| pCiMX254 | *M. smegmatis* | pCi46 (*nuoD*_a) | *qcrB*_b | pCi51 | MMO120 | MMO121 |
| pCiMX255 | *M. smegmatis* | pCi46 (*nuoD*_a) | *ctaC*_b | pCi53 | MMO120 | MMO121 |
| pCiMX256 | *M. smegmatis* | pCi46 (*nuoD*_a) | *atpE*_a | pCi54 | MMO120 | MMO121 |
| pCiMX257 | *M. smegmatis* | pCi46 (*nuoD*_a) | *menD*_b | pCi68 | MMO120 | MMO121 |
| pCiMX258 | *M. smegmatis* | pCi46 (*nuoD*_a) | *mq*o_b | pCi60 | MMO120 | MMO121 |
| pCiMX259 | *M. smegmatis* | pCi46 (*nuoD*_a) | *sdhA1*_a | pCi3 | MMO120 | MMO121 |
| pCiMX260 | *M. smegmatis* | pCi46 (*nuoD*_a) | *cydB*_b | pCi66 | MMO120 | MMO121 |
| pCiMX262 | *M. smegmatis* | pCi4 (*sdhA2*_a) | *ndh*_b_3MM | pCi267 | MMO120 | MMO121 |
| pCiMX263 | *M. smegmatis* | pCi51 (*qcrB*_b) | *ndh*_b_3MM | pCi267 | MMO120 | MMO121 |
| pCiMX264 | *M. smegmatis* | pCi53(*ctaC*_b) | *ndh*_b_3MM | pCi267 | MMO120 | MMO121 |
| pCiMX265 | *M. smegmatis* | pCi54 (*atpE*_a) | *ndh*_b_3MM | pCi267 | MMO120 | MMO121 |
| pCiMX266 | *M. smegmatis* | pCi68(*menD*_b) | *ndh*_b_3MM | pCi267 | MMO120 | MMO121 |
| pCiMX267 | *M. smegmatis* | pCi60(*mqo*_b) | *ndh*_b_3MM | pCi267 | MMO120 | MMO121 |
| pCiMX268 | *M. smegmatis* | pCi3 (*sdhA1*) | *ndh*_b_3MM | pCi267 | MMO120 | MMO121 |
| pCiMX269 | *M. smegmatis* | pCi66 (*cydB*_b) | *ndh*_b_3MM | pCi267 | MMO120 | MMO121 |
| pCiMX271 | *M. smegmatis* | pCi4 (*sdhA2*_a) | *ndh*_b_5MM | pCi268 | MMO120 | MMO121 |
| pCiMX272 | *M. smegmatis* | pCi51 (*qcrB*_b) | *ndh*_b_5MM | pCi268 | MMO120 | MMO121 |
| pCiMX273 | *M. smegmatis* | pCi53(*ctaC*_b) | *ndh*_b_5MM | pCi268 | MMO120 | MMO121 |
| pCiMX274 | *M. smegmatis* | pCi54 (*atpE*_a) | *ndh*_b_5MM | pCi268 | MMO120 | MMO121 |
| pCiMX275 | *M. smegmatis* | pCi68(*menD*_b) | *ndh*_b_5MM | pCi268 | MMO120 | MMO121 |
| pCiMX276 | *M. smegmatis* | pCi60(*mqo*_b) | *ndh*_b_5MM | pCi268 | MMO120 | MMO121 |
| pCiMX277 | *M. smegmatis* | pCi3 (*sdhA1*) | *ndh*_b_5MM | pCi268 | MMO120 | MMO121 |
| pCiMX278 | *M. smegmatis* | pCi66 (*cydB*_b) | *ndh*_b_5MM | pCi268 | MMO120 | MMO121 |
| pCiMX279 | *M. smegmatis* | pCi49 (*ndh*_b) | *atpE*_a_3MM | pCi285 | MMO120 | MMO121 |
| pCiMX280 | *M. smegmatis* | pCi4 (*sdhA2*_a) | *atpE*_a_3MM | pCi285 | MMO120 | MMO121 |
| pCiMX281 | *M. smegmatis* | pCi51 (*qcrB*_b) | *atpE*_a_3MM | pCi285 | MMO120 | MMO121 |
| pCiMX282 | *M. smegmatis* | pCi53 (*ctaC*_b) | *atpE*_a_3MM | pCi285 | MMO120 | MMO121 |
| pCiMX284 | *M. smegmatis* | pCi68 (*menD*_b) | *atpE*_a_3MM | pCi285 | MMO120 | MMO121 |
| pCiMX285 | *M. smegmatis* | pCi60 (*mqo*_b) | *atpE*_a_3MM | pCi285 | MMO120 | MMO121 |
| pCiMX286 | *M. smegmatis* | pCi3 (*sdhA1*) | *atpE*_a_3MM | pCi285 | MMO120 | MMO121 |
| pCiMX287 | *M. smegmatis* | pCi66 (*cydB*_b) | *atpE*_a_3MM | pCi285 | MMO120 | MMO121 |
| pCiMX288 | *M. smegmatis* | pCi285 (*atpE*_a_3MM) | *ndh*_b_3MM | pCi267 | MMO120 | MMO121 |
| pCiMX289 | *M. smegmatis* | pCi285 (*atpE*_a_3MM) | *ndh*_b_5MM | pCi268 | MMO120 | MMO121 |
